# Phosphorylation of the RB C-terminus regulates condensin II release from chromatin

**DOI:** 10.1101/2020.10.22.350199

**Authors:** Seung J. Kim, James I. MacDonald, Frederick A. Dick

## Abstract

The retinoblastoma tumour suppressor protein (RB) plays an important role in biological processes such as cell cycle control, DNA damage repair, epigenetic regulation, and genome stability. The canonical model of RB regulation is that cyclin-CDKs phosphorylate, and render RB inactive in late G1/S, promoting entry into S phase. Recently, mono-phosphorylated RB species were described to have distinct cell-cycle independent functions, suggesting that a phosphorylation code dictates diversity of RB function. However, a biologically relevant, functional role of RB phosphorylation at non-CDK sites has remained elusive. Here, we investigated S838/T841 dual phosphorylation, its upstream stimulus, and downstream functional output. We found that mimicking T-cell receptor activation in Jurkat leukemia cells induced sequential activation of downstream kinases including p38 MAPK, and RB S838/T841 phosphorylation. This signaling pathway disrupts RB and condensin II interaction with chromatin. Using cells expressing a WT or S838A/T841A mutant RB fragment, we present evidence that deficiency for this phosphorylation event prevents condensin II release from chromatin.

## INTRODUCTION

The retinoblastoma tumour suppressor protein (RB) is well-characterized for its role in regulating entry into the cell cycle, and this is its canonical function. In G1, RB is bound to E2F transcription factors, repressing their ability to express cell cycle target genes (1, 2). Upon growth stimulation, cyclin D accumulates and binds to cyclin-dependent-kinases 4/6 (CDK4/6). Cyclin D-CDK4/6 complexes phosphorylate RB, partially relieving E2F repression (3). This leads to expression of cyclin E, and cyclin E-bound CDK2 hyperphosphorlyates RB. Hyperphosphorylated RB is thought to be inactivated, and free E2F transcription factors drive gene expression required for subsequent cell cycle progression (4–6).

This model, however, represents only one facet of RB function, and CDK-independent roles are emerging. These non-canonical functions are ascribed to a pool of chromatin-bound RB that localize to various parts of the genome. For example, RB is required for recruiting DNA repair machinery for non-homologous end joining (NHEJ) and homology-directed repair (HR) pathways at double strand breaks (DSBs) (7, 8). In addition, RB recruits epigenetic writers to repetitive elements that deposit repressive histone marks (9, 10). Heterochromatinization of repetitive elements may be required to silence their transcription to avoid an autoimmune response, and maintain chromosome stability (10–13). In particular, RB-dependent maintenance of histone 4 lysine 20 tri-methylation (H4K20me3) at pericentromeric repeats is required for proper chromatid segregation in anaphase (10). Furthermore, disrupting RB interaction with structural maintenance of chromatin containing complexes such as condensin II at pericentromeric repeats results in mitotic errors, replication defects and aneuploidy (14, 15). Therefore, existing literature suggests that there is a pool of RB that is resistant to CDK inactivation and binds chromatin to carry out cell cycle-independent functions.

Mechanistically, post-translational modifications regulate RB’s diverse functions (16). Notably, phosphorylation of RB can regulate its cell cycle-independent functions. For example, CDK4/6 phosphorylation of RB at S249/T252 allows RB to bind and inactivate the NFκB family protein p65 and its immune-suppressive transcriptional program in prostate cancer cells (17). In addition, stress-induced p38 mitogen activated protein kinase (MAPK) activation phosphorylates RB at S249/T252 to negate inactivation by CDKs and prevent cell cycle entry (18). Furthermore, mono-phosphorylation of RB at its thirteen CDK phosphorylation sites confer ability to activate distinct transcriptional programs beyond cell cycle genes such as inflammatory response and mitochondrial oxidative phosphorylation (19). However, these studies have primarily focused on phosphorylation at CDK consensus sites, and RB phosphorylation at non-CDK sites and their functional outcomes are not fully characterized. Upon apoptotic stimulation, p38 MAPK, but not CDKs, phosphorylate RB at non-CDK sites, and activates E2F1 transcription in a reporter assay (20). Specific non-CDK phosphorylation sites, and their relevance to cellular function remain to be elucidated.

Here, we characterize RB phosphorylation on S838 and T841 (S838/T841) as a mechanistic link between T-cell receptor (TCR) activation and chromatin decondensation in Jurkat T-cells. Prompted by existing phospho-proteomic data, we generated an antibody specific to phosphorylation at S838/T841. We show that p38 MAPK is an upstream kinase that induces RB phosphorylation at these sites upon stimulus that mimics TCR activation in culture. Finally, we demonstrate that inability to phosphorylate these sites leads to defective condensin II release from chromatin.

## RESULTS

### Detection of C-terminal RB phosphorylation on S838 and T841

Multiple residues located throughout RB are phosphorylated, and the functional roles of CDK sites have been extensively characterized. Many of these modifications serve to inactivate its canonical function of negatively regulating cell cycle entry through E2F transcription factor binding. Interestingly, RB mono-phosphorylation of CDK sites has been shown to activate specific molecular functions of RB that may extend beyond cell cycle control (19). We sought to identify RB phosphorylation at new sites and determining their roles in RB regulation.

Figure 1A depicts the open reading frame of RB, its known CDK phosphorylation sites (top) and potential phospho-acceptor residues that are uncharacterized (bottom). We focused on RB C-terminus (RBC) as it forms an α-helix, and any phosphorylation within this region is likely to disrupt protein-protein interactions, leading to a novel functional outcome. To identify candidate phosphorylation sites for our investigation, we curated predicted phosphorylation events within tryptic fragments corresponding to RBC from available phospho-proteomic data in different cells or tissues upon various treatments (Figure 1B) (21–29). A positive prediction for each serine or threonine residue is shaded in blue.

**Figure 1:**
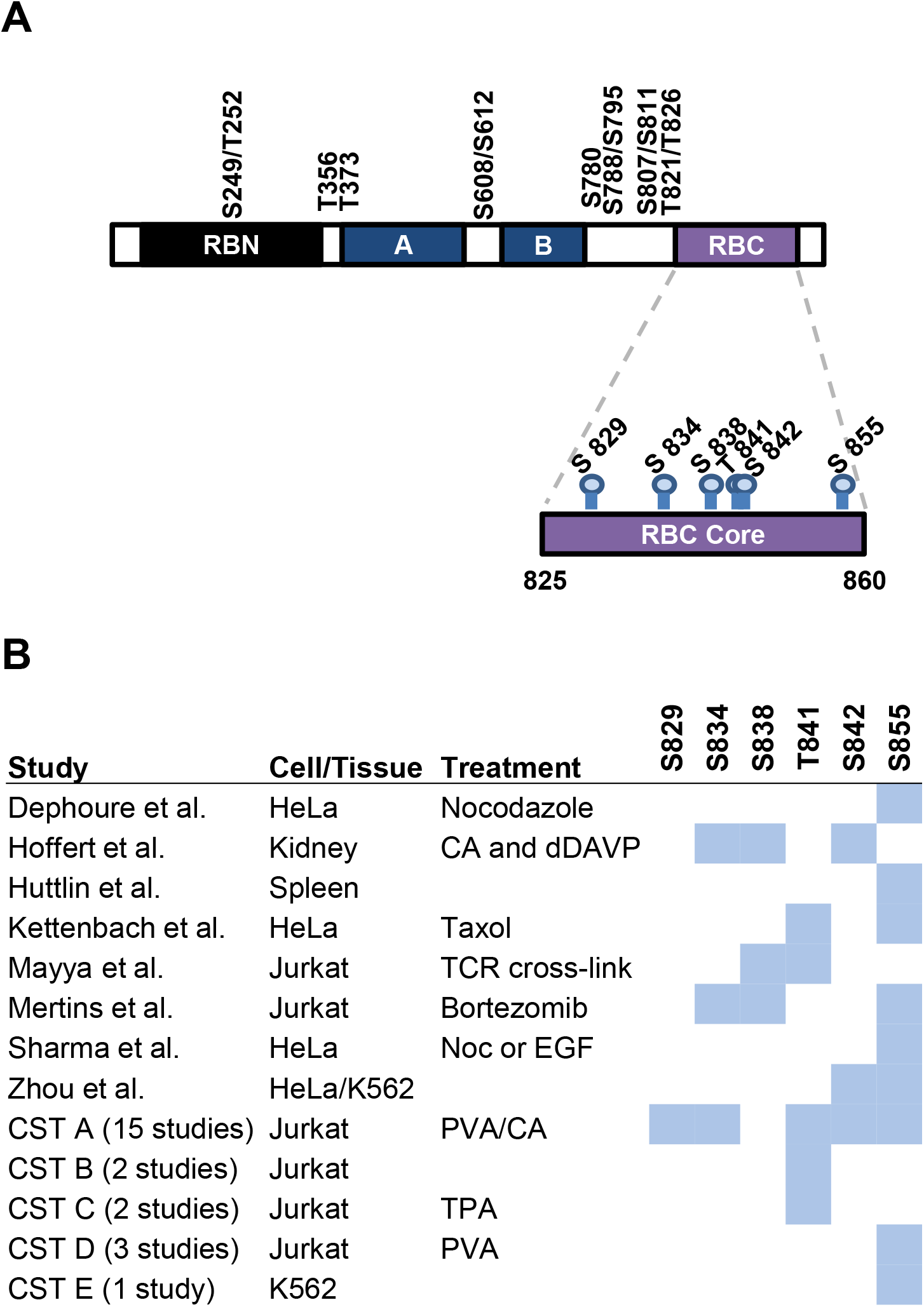
Putative phosphorylation of the RB C-terminus. **(A)** The open reading frame for RB is shown depicting known CDK phosphorylation sites (top), and uncharacterized, non-CDK phosphorylation sites in RBC (bottom). RB N-terminal domain (RBN), A and B halves of the pocket domain, and RB C-terminal domain (RBC) are shown. **(B)** Chart displaying potential phosphorylation sites in the amino acid 825-860 region of the RB C-terminus. Putative phosphorylation sites identified in publicly available phosphoproteomic studies are shaded in blue and the relevant data sources are indicated. CST – Cell Signalling Technologies data set.

The phospho-proteomic data reveal various combinations of serine and threonine residues within RBC are readily phosphorylated, supporting the idea that the RB phosphorylation “code” is yet incomplete. There is significant variability between the data sets on which exact sites are phosphorylated. This may indicate that complex, differential phosphorylation events occur in RBC depending on the cell type and stimulus, or that whole cell proteomics cannot accurately discern between neighbouring phosphorylation events. While all of these potential modifications can be found in untreated cells, we focused on S838 and T841 phosphorylation because it is inducible upon TCR activation, a relevant signaling pathway in Jurkat leukemic T-cells.

In order to study S838 and T841 phosphorylation on RB further, we sought to generate a specific antibody for these modifications. We generated rabbit anti serum against an RB derived peptide corresponding to amino acids 834-845, phosphorylated on S838 and T841. An ELISA was performed to confirm the specificity of our antibodies following purification from the anti-serum. We found that anti-RB pS838/pT841antibodies had a 10-fold higher affinity for the S838/T841 phosphorylated RB peptide than its unphosphorylated counterpart (Figure 2A). Next, we sought to find evidence for RB phosphorylation on S838/T841 *in vitro* as predicted by phospho-proteomic data. We incubated glutathione S-transferase-tagged RBC (GST-RBC) with whole cell extract (Ext) from Jurkat cells treated with phosphatase inhibitors pervanadate and calyculin A (PVA/CA) to globally activate kinases that may phosphorylate RB. We found a time-dependent increase in RB S838/T841 phosphorylation in the presence of ATP (Figure 2B). To account for the possibility that our antibody cross-reacts with other phosphorylation events in RBC (amino acids 792-928), we repeated the experiment with GST-RBC where S838 and T841 were substituted to alanine either together or as single mutants. This reveals that this antibody requires either S838 or T841 phosphorylation, but not both (Figure 2C). For this reason, we describe the antibodies and detected phosphorylation events throughout this paper as pS838/pT841 because phosphorylation of either site alone, or both in combination, is possible. With the specificity of our antibody established, we determined if endogenous RB is phosphorylated on S838/T841. We treated Jurkat cells with PVA/CA to activate kinases and RB phosphorylation, prepared nuclear extracts (NE) and immunoblotted with our antibody. RB phosphorylation on S838/T841 was increased upon PVA/CA treatment, but it was also evident that our antibody cross-reacted with approximately six or seven proteins other than RB (Figure 2D). To ensure we were certain of RB’s identity, we repeated the cell treatment and immunoprecipitated total RB from Jurkat NE and performed an immunoblot with anti-RB pS838/pT841antibodies. Once again, we confirmed that RB phosphorylation on S838/T841 was induced upon PVA/CA treatment (Figure 2E). In all subsequent experiments, we first immunoprecipitated RB then probed with the antibody to assess S838/T841 phosphorylation.

**Figure 2:**
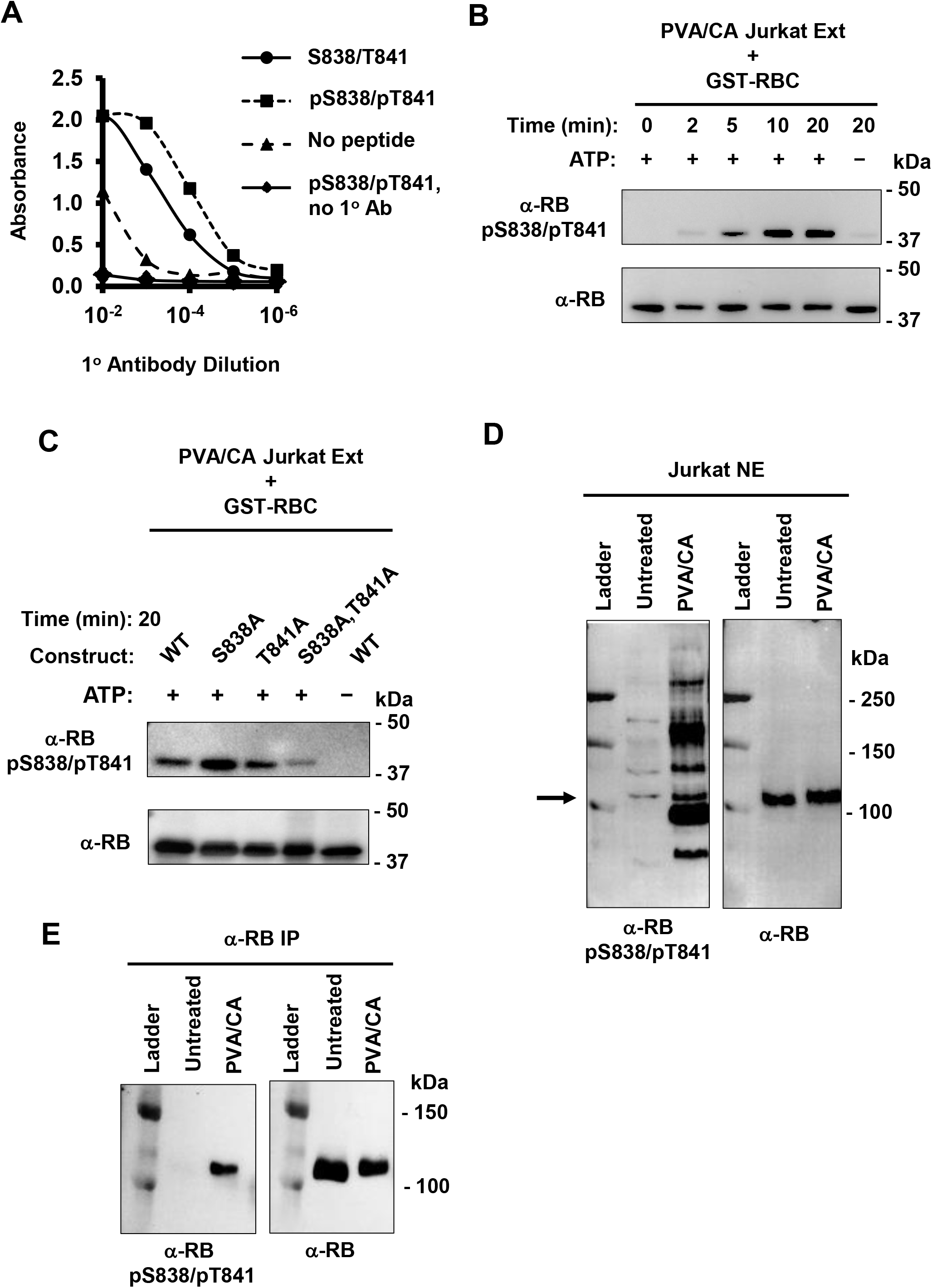
Detection of phosphorylation of S838/T841 on RB. **(A)** Rabbit anti-serum was generated by immunizing animals with a peptide phosphorylated at S838 and T841. Phosphorylation-specific antibodies were purified and an ELISA was performed to determine their specificity. Phosphorylated or unphosphorylated peptides were adhered to microplate wells and incubated with anti-RB pS838/pT841antibodies. Control wells contained either no peptide or no antibody. Antibody binding was measured colorimetrically and graphed. **(B)** A GST-tagged RBC fragment (aa 792-928) was incubated with PVA/CA treated Jurkat whole cell extract for the indicated times, and omission of ATP was used as a negative control. Phosphorylation status was determined by immunoblotting with the anti-RB pS838/pT841, and blotting with anti-RB antibodies was used as a loading control. **(C)** WT or mutant GST-RBC proteins were incubated with PVA/CA treated Jurkat extracts with or without ATP for 20 min. Phosphorylation status was measured as in B. **(D)** S838/T841 phosphorylation of endogenous RB was detected by western blotting with anti-RB pS838/pT841 antibodies in PVA/CA treated Jurkat nuclear extracts (NE). This membrane was stripped and reprobed for RB and the arrow indicates the corresponding RB band. **(E)** Jurkat cells were treated with PVA/CA, and RB was immunoprecipitated from NE with an α-RB antibody-coupled to Protein G Dynabeads. Phosphorylation status of S838/T841 was determined by western blotting.

These data demonstrate that our antibody is highly specific for phosphorylation within RBC, namely on S838/T841. Although it cross-reacts with a small number of nuclear proteins, first enriching for RB by immunoprecipitation is a robust and specific means of detecting phosphorylation of these sites. More importantly, we have shown the first line of evidence that S838/T841 phosphorylation is a bona fide RB post-translational modification that has not yet been studied functionally.

### Sequential activation of kinases in the TCR signalling pathway induces RB S838/T841 phosphorylation

As a first step to understanding the functional role for RB S838/T841 phosphorylation, we sought to identify the kinase that was responsible for this modification. p38 MAPK has been shown to phosphorylate RB even when CDK sites are mutated (20). Therefore, we pre-treated Jurkat cells with a p38 inhibitor, SB203580, then stimulated phosphorylation with PVA/CA. We found that SB203580 pre-treatment strongly reduced RB S838/T841 phosphorylation despite PVA/CA treatment (Figure 3A). As a positive control for p38 inhibition, we also measured phosphorylation of a known p38 target, MAPKAPK-2, and found that phosphorylation of its activation loop at T334 was also reduced. Next, we determined if immunoprecipitated p38 can phosphorylate RB at S838/T841 *in vitro*. We first isolated p38 by immunoprecipitating the active form with anti-p38 pT180/pY182 phosphorylation specific antibodies. It was then incubated with GST-RBC in the presence or absence of its inhibitor. p38 phosphorylation of RB increased over time, and its inhibitor largely abrogated this effect (Figure 3B).

**Figure 3:**
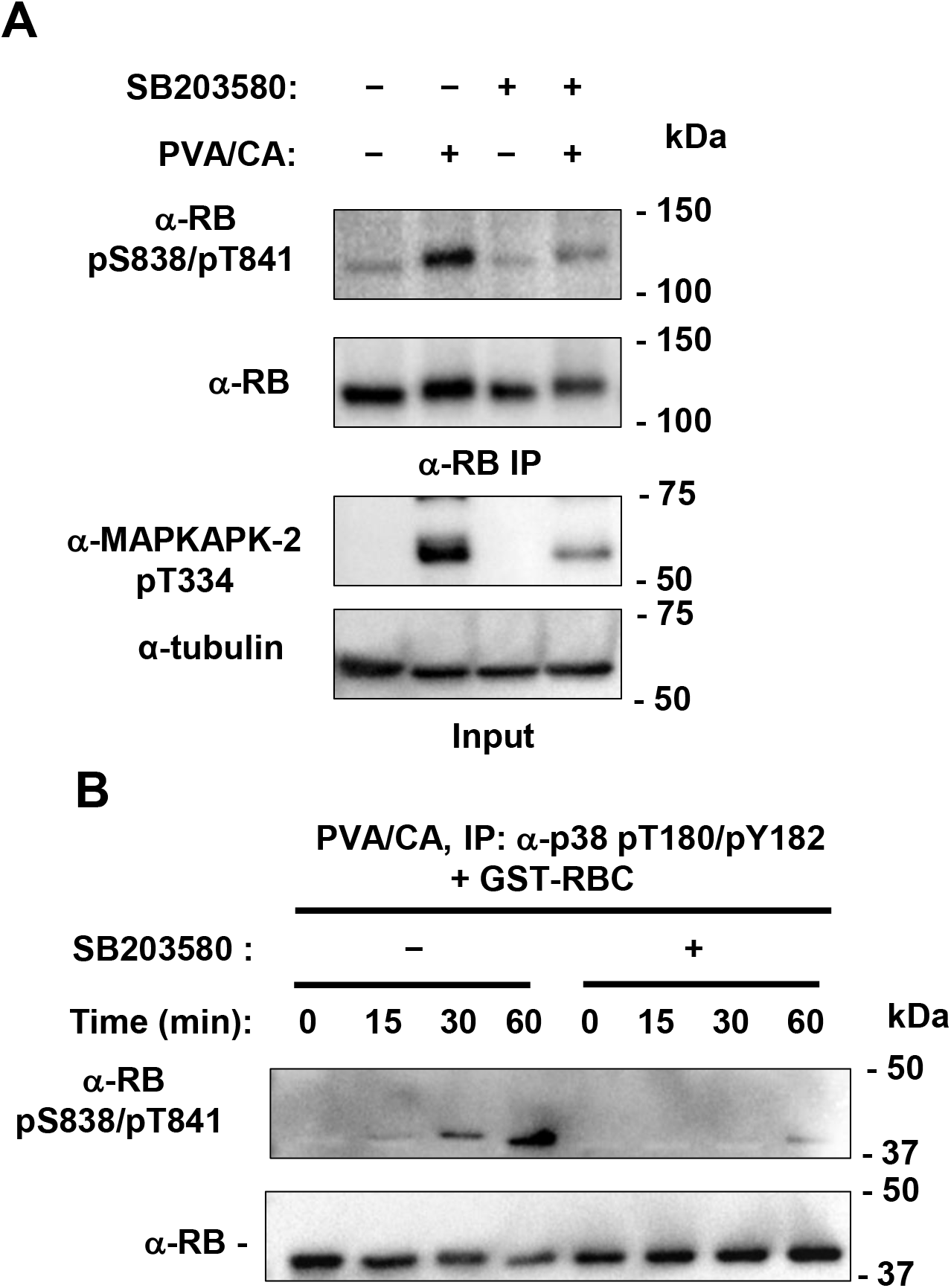
Phosphorylation of RB on S838/T841 is dependent on p38. **(A)** Jurkat cells were treated with PVA/CA with or without 10 μM SB203580, a p38 inhibitor. RB was immunoprecipitated from lysates with an α-RB antibody bound to Protein G Dynabeads, and phosphorylation of S838/T841 was detected by immunoblotting. As a positive control for p38 inhibition, phosphorylation of a known p38 phosphorylation target, T334 on MAPKAPK-2, was detected by western blotting. **(B)** Jurkat cells were treated with PVA/CA, and active p38 was immunoprecipitated from whole cell lysates with α-p38 pT180/pY182 antibodies bound to Protein G Dynabeads. Immunoprecipitated p38 was resuspended in kinase buffer containing GST-RBC, with or without 50 μM SB203580 for the indicated amount of time. Phosphorylation of S838/T841 and GST-RBC was detected by western blotting as indicated.

These experiments strongly suggest that RB phosphorylation at S838/T841 is dependent on p38. PVA/CA treatment indiscriminately activates kinases, yet selective inhibition of p38 was sufficient to reduce RB phosphorylation to background level. Furthermore, partially purified, active p38 was necessary and sufficient to induce RB phosphorylation *in vitro* suggesting it may do so directly.

PVA/CA treatment has served as a way to easily and efficiently induce phosphorylation of RB. However, this treatment activates many pathways and hence does not offer insight into specific stimuli that induce RB S838/T841 phosphorylation under physiological conditions. Since p38 is a downstream target of T-cell receptor (TCR) signalling, and Jurkat cells are leukemic T-cells, we hypothesized that TCR activation phosphorylates RB through p38. We mimicked TCR activation by treating cells with an antibody cocktail that physically aggregates TCR and its co-stimulatory receptor together, as previously reported (30, 31). We then detected phosphorylation-dependent activation of proteins in the signalling pathway by immunoblotting. We observed rapid activation of ZAP70 within the first five minutes of antibody crosslinking, and peak activation of p38 soon after at 15 minutes (Figure 4A). TCR activation reproducibly induced p38 activation within the first 30 minutes (Figure 4B). At 30 minutes we immunoprecipitated RB and found strong S838/T841 phosphorylation (Figure 4C). In addition, TCR activation did not induce additional phosphorylation of neighbouring CDK phosphorylation sites, S807 and S811 consistent with these cells already being in a proliferative state. S838/T841 phosphorylated RB also retained its interaction with E2F1 and E2F2 as they were comparably co-immunoprecipitated relative to untreated cells. The shift in E2F2 migration upon TCR activation is consistent with previous proteomic data that report S117/S123 phosphorylation in Jurkat cells upon PVA/CA or TCR activation (25, 29).

**Figure 4:**
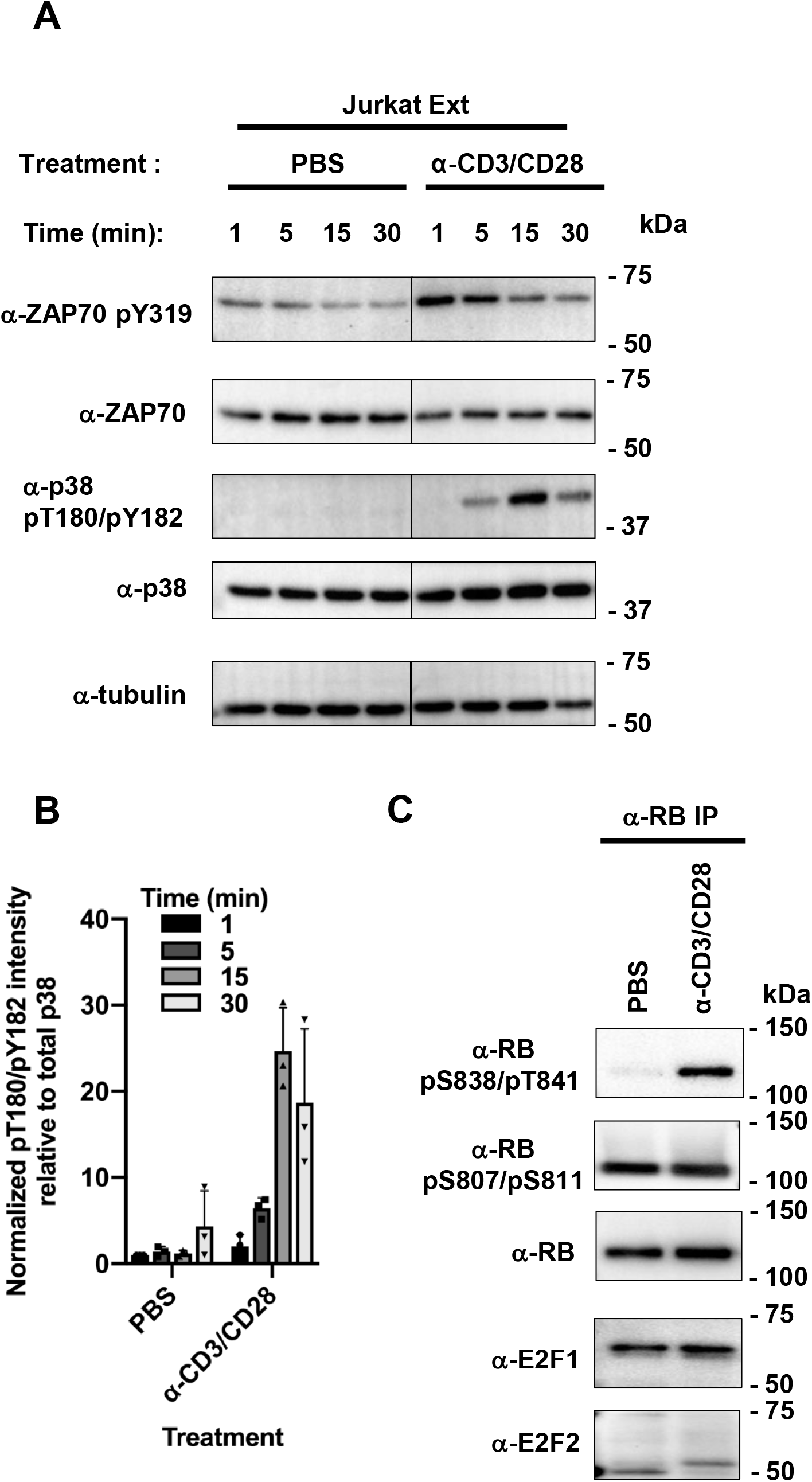
T-cell receptor signalling causes RB S838/T841 phosphorylation. **(A)** Jurkat cells were treated with PBS, or anti-CD3, anti-CD28, and a secondary antibody to cross link T-cell receptors for the indicated time points. Each time point was quenched by washing cells with cold PBS and immediately lysed to generate whole cell extracts. Abundance of phosphorylated species relative to total ZAP70 and p38 was determined with immunoblotting as a read-out. Relevant lanes from a single whole image was spliced together at the vertical line in the center for each blot. **(B)** Jurkat cells were treated as in A for 30 minutes. RB was immunoprecipitated from whole cell extracts with α-RB antibodies bound to Protein G Dynabeads. RB phosphorylation on S838/T841 and S807/S811 was determined by blotting, and co-immunoprecipitating E2Fs were detected similarly.

These experiments implicate TCR activation as a relevant mechanism that induces RB S838/T841 phosphorylation and suggests that RB S838/T841 phosphorylation may regulate its non-canonical functions. Our antibody cocktail treatment appears to closely mimic TCR activation as both ZAP70 and p38 were activated sequentially in the anticipated order. A slight temporal delay in peak phosphorylation of ZAP70, p38 and ultimately RB also suggests that this signalling cascade is biologically relevant to TCR activation. Furthermore, because S838/T841 phosphorylation in response to TCR signalling is unlinked to CDK phosphorylation its function is unlikely to be related to cell cycle entry or control of E2F-dependent transcription.

### RB S838/T841 phospho-acceptor mutants prevent condensin II unloading and chromatin decondensation

The majority of previously described RB function stems from its ability to occupy chromatin. For example, RB protects genome integrity during mitosis by recruiting the condensin II complex to pericentromeric heterochromatin (15). In addition, the condensin II complex has a role in developing T-cells. The *nessy* mutant allele of CAPH2, one of the subunits of the complex, results in defective condensation of T-cell chromatin and development (32, 33). Since RB is phosphorylated under conditions that mimic TCR signalling, and is known to recruit condensin II to chromatin, we hypothesized that RB S838/T841 phosphorylation may regulate RB-condensin II interactions on chromatin in T-cells.

In order to compare chromatin occupancy upon TCR crosslinking, we first sought to establish that this stimulus did not affect total expression of the proteins of interest. Indeed, we confirmed that expression of CAPH2, RB, E2F1 and SMC1, a cohesin subunit, was not changed in whole cell extracts (Figure 5A, left). However, TCR crosslinking for 30 minutes induced release of RB, E2F1, and CAPH2, but not SMC1 from chromatin (Figure 5A, right). Using densitometric analysis of these experiments we found that TCR crosslinking released 40% of CAPH2 from chromatin (Figure 5B). To examine the precise role of S838/T841 phosphorylation in this process, we created a phospho-acceptor mutation in RB with a S838A/T841A double alanine substitution. To stably express this mutant in cells, we generated lentiviral vectors coding for HA-tagged large pocket fragment of RB (RBLP, amino acids 379-928). Jurkat cells were transduced with lentiviruses expressing either the WT RBLP (RBLP-WT) or S838A/T841A (RBLP-AA) construct (Figure 5C). We confirmed equivalent expression of both constructs by immunoblotting for the HA-tag (Figure 5D). We cross-linked TCRs as before and investigated CAPH2 and SMC1 chromatin levels (Figure 5E). This revealed that RBLP-AA expressing cells retained similar levels of chromatin-bound CAPH2 as unstimulated cells and that RBLP-WT expressing cells released CAPH2 similarly to untransduced Jurkat cells (Figure 5E). We normalized CAPH2 release to SMC1 bound chromatin levels to quantitate the inhibition of CAPH2 release by RBLP-AA (Figure 5F). These experiments confirm that the inability to phosphorylate RB at S838/T841 dominantly blocks condensin II release from chromatin.

**Figure 5:**
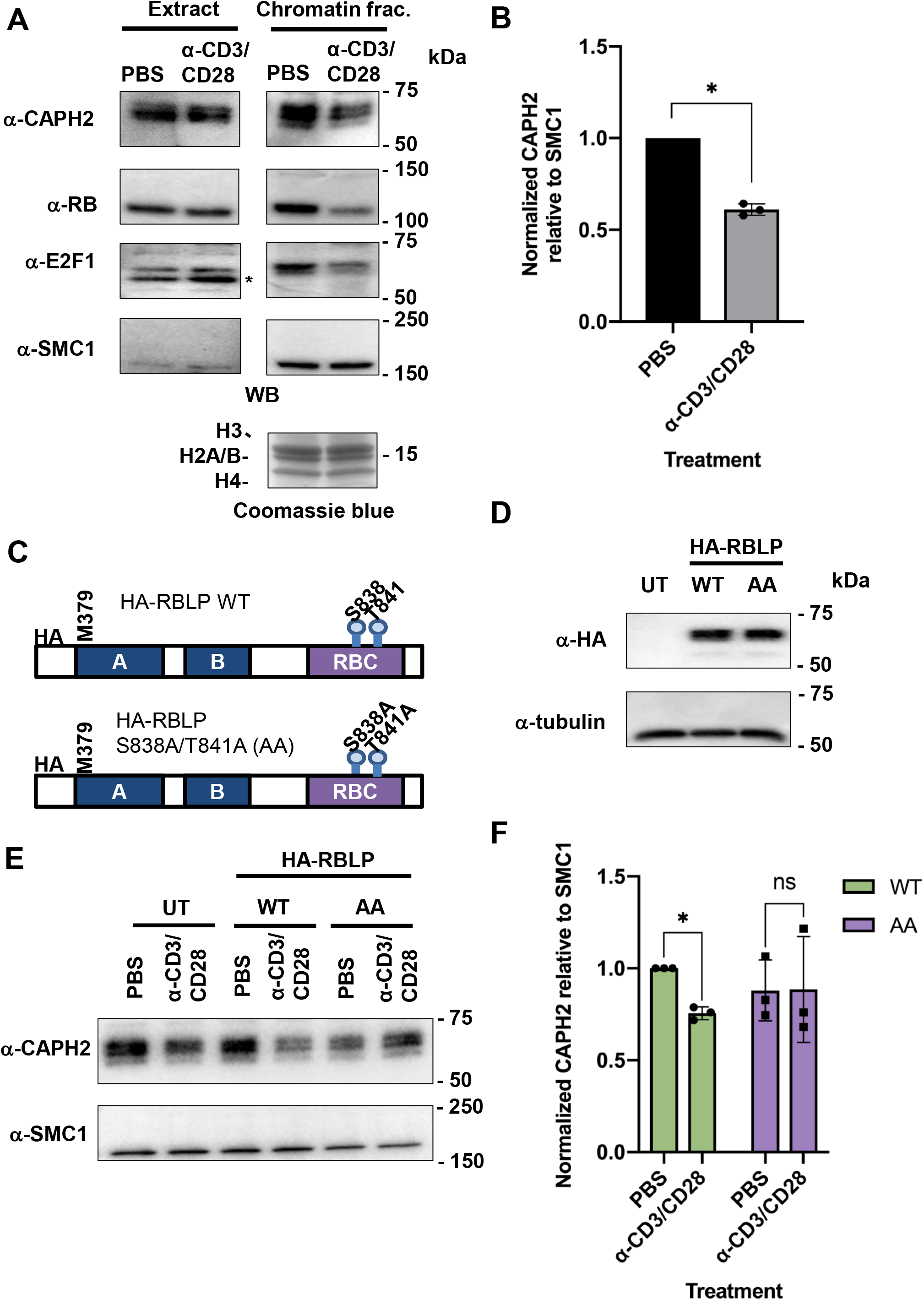
Non-phosphorylatable S838A/T841A RB prevents CAPH2 chromatin unloading upon T-cell receptor crosslinking. **(A)** Jurkat cells were treated as indicated for 30 minutes, whole cell extracts and chromatin fractions were prepared, and expression levels of the indicated proteins were detected by western blotting. Histone levels serve as loading control and were detected by Coomassie Blue staining. * indicates non-specific band. **(B)** Relative levels of CAPH2 in chromatin fractions was normalized to SMC1 and compared between treated and untreated conditions (n=3, * *P* < 0.05, Student’s t-test). **(C)** The open reading frame for WT and mutant RBLP constructs (aa 379-928) are shown depicting their relevant coding regions and HA tags. The mutant construct labelled RBLP AA carries double alanine substitutions at S838 and T841. **(D)** Lentiviruses were used to transduce RBLP and the AA mutant construct into Jurkat cells. Expression of exogenous RBLP was detected by western blotting for HA. Tubulin blots serve as loading controls. **(E)** Jurkat cells expressing either WT or RBLP, or untransduced controls (UT) were subjected to TCR crosslinking for 30 minutes. Chromatin fractions were prepared, and levels of the indicated proteins were detected by western blotting. **(F)** Levels of chromatin associated CAPH2 was normalized to SMC1 and compared between PBS and TCR crosslinked conditions (n=3, * *P* < 0.05, Two-way ANOVA and Sidak’s multiple comparisons test).

Next, we sought to describe a biochemical consequence of defective CAPH2 unloading. To assess changes in chromatin structure upon TCR-crosslinking, we used sensitivity to sonication-induced chromatin shearing as a read-out for chromatin compaction as previously reported (Figure 6A)(33). Untreated or TCR crosslinked cells were fixed to immobilize DNA-protein contacts and fractionated to obtain total chromatin. Isolated chromatin was sheared by a range of sonication cycles, and the presence of high molecular weight DNA fragments was detected by agarose gel electrophoresis and ethidium bromide staining. In line with Figure 5A, TCR crosslinking decreased the amount of high molecular weight DNA remaining after the same number of sonication cycles compared to control cells that were untreated (Figure 6B-C). We repeated this TCR crosslinking experiment using cells that express RBLP-WT or the - AA mutant. This revealed that chromatin isolated from RBLP-AA expressing cells was sheared less effectively compared to that from RBLP-WT expressing cells (Figure 6D-E). This suggests that the RB S838A/T841A mutant disrupts TCR crosslink-mediated chromatin decondensation and suggests that the phosphorylation of RB on these sites plays a key role in chromatin decondensation by releasing RB and condensin II from chromatin.

**Figure 6:**
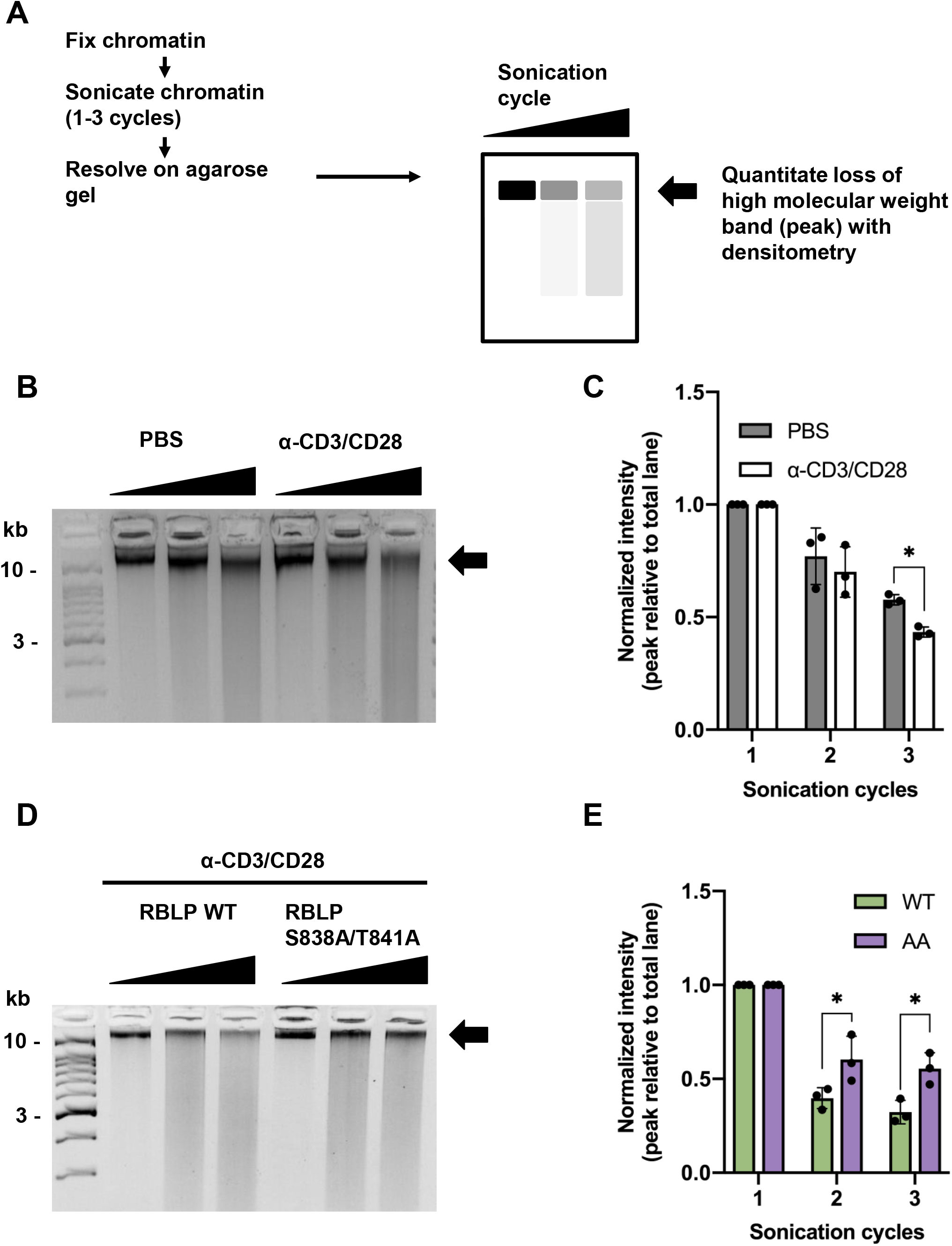
RB S838/T841 phosphorylation regulates chromatin dynamics upon T-cell receptor signaling. **(A)** Jurkat cells were stimulated by T-cell receptor crosslinking as indicated. Cells were fixed and lysed to obtain chromatin fractions. Chromatin was sonicated for one to three cycles. Protein-DNA crosslinks were reversed and DNA was purified, and analyzed on a 3% agarose gel. DNA was stained with ethidium bromide and visualized on a Chemi Doc. The high molecular weight DNA band indicated by the arrow was quantified on Image Lab and intensity is expressed as a percentage of the total lane normalized within in each group. **(B)** Untransduced Jurkat cells were PBS or TCR crosslinked as described in A. Chromatin fragmentation is shown. **(C)** Percent high molecular weight DNA quantities were averaged and graphed (n = 3, * indicates *P* < 0.05, two-way ANOVA with Sidak’s multiple comparisons test). **(D)** Lentiviral transduced Jurkat cells were stimulated by T-cell receptor crosslinking, fixed, and lysed to obtain chromatin fractions. Chromatin was sonicated for increasing number of cycles and analyzed as in A. **(E)** The high molecular weight band indicated by the arrow was quantified, averaged and graphed, (n = 3, * indicates *P* < 0.05, Two-way ANOVA with Sidak’s multiple comparisons test).

## DISCUSSION

In this report, we characterized a functional role for non-CDK phosphorylation of RB. By curating existing phospho-proteomic data, we generated and validated S838/T841 phospho-specific antibodies. With this tool, we showed that p38 MAPK is an upstream kinase that can direct RB phosphorylation, and that TCR crosslinking induces a signaling cascade that ultimately phosphorylates RB in Jurkat cells on these sites. To probe its mechanistic role, we generated cell lines that stably express phospho-acceptor mutant RB. Finally, our data demonstrates that when RB is unable to be phosphorylated at S838/T841, TCR signaling is unable to release condensin II from chromatin. This conclusion is consistent with previous work that describes RB-condensin II chromatin interactions and T-cell development, and contributes to our understanding of an emerging RB phosphorylation code.

TCR crosslinking induced marked loss of chromatin bound CAP-H2, but not SMC1, suggesting large scale, yet selective reorganization of chromatin architecture. Importantly, expression of RBLP-AA was sufficient to disrupt TCR crosslinking induced CAP-H2 release. The selective nature of RB regulation of condensin II in Jurkat T-cells agrees with the previous characterization of the *nessy* mutant CAP-H2 allele that is responsible for defective T-cell differentiation. Recently published work may inform future studies on specific consequences of the release of CAP-H2 in this model. CAP-H2 and transcription factor IIIC (TFIIIC) localize to histone clusters in mouse embryonic stem cells, and serve to maintain topologically-associated-domains that regulates transcription (34). In addition, RB-condensin II complexes can regulate transcription at bi-directional promoters, and maintain long-range chromosomal interactions. ChIP-sequencing (ChIP-Seq) for condensin II and TFIIIC components upon TCR crosslinking may inform site-specific loss of interaction, and the mechanistic role of RB can be delineated with a phospho-acceptor RB mutant. For increased sensitivity, gene editing technologies should be adopted to mutate endogenous RB, or deplete endogenous RB with an inducible system as previously reported. ChIP-seq data should be interpreted together with high-throughput chromosome conformation capture (3C) such as 4C-seq to fully characterize functional consequences.

We speculate that RB phosphorylation at S838/T841 may be a key step in naive T-cell activation. Previous work has demonstrated that chromatin condensation orchestrated by condensin II is required to avoid cell death in developing thymocytes (33). Once mature, these cells become quiescent. Upon TCR activation through cognate antigen binding and co-stimulation, re-entry into cell cycle is coupled with chromatin decondensation and nuclear expansion in naïve T-cells. Loss of CAP-H2 chromatin occupancy upon TCR crosslinking in Jurkat T-cells is in line with such macromolecular changes during T-cell activation. Furthermore, TCR crosslinking with CD3/CD28 antibodies specifically induced RB phosphorylation at S838/T841, but not at its CDK sites. RB interaction with E2F transcription factors was also unaffected. These data suggest that S838/T841 phosphorylation is functionally distinct from canonical cell-cycle control by RB that may also be cell type specific. Future work with peripheral T-cells *ex vivo* is warranted to support this model.

Ultimately, our work adds a layer of complexity to phosphorylation dependent regulation of RB function. The canonical model of cell cycle control by RB has been binary; cyclin-CDK driven RB hyperphosphorylation renders RB’s negative regulation of E2F transcription factors inactive. Recently, however, various mono-phosphorylated RB species at one of its CDK sites have been shown to regulate cell-cycle independent transcription programs. We showed that cell-cycle independent TCR crosslinking induces a signaling cascade via p38 that can ultimately phosphorylate RB at S838/T841. Future work is warranted to characterize the relevance and functional roles of S838/T841 phosphorylation in other physiological circumstances.

## EXPERIMENTAL PROCEDURES

### Antibodies

Anti-RB pS838/pT841 antibody was produced in rabbits by Covance (Pennsylvania, USA). The peptide immunogen was corresponded to the sequence 834-SIGE(pS)FG(pT)SEKF-845 of pRB. Peptides were immobilized using the Sulfo-link coupling reagent (Thermo-Fisher) following the manufacturer’s recommendations. The phospho-specific antibody was purified using standard procedures by passing serum first over the non-phosphorylated peptide column and secondly over the phosphorylated-peptide column to remove any antibodies recognizing the non-phosphorylated species. Anti-RB pS838/pT841 antibody was used in primary antibody solution at 0.5 mg/mL for Western blotting. Anti-MAPKAPK-2 pT334 (27B7), p38 pT180/pY182 (9211), total p38 (9212), α-tubulin (11H10) and RB pS807/pS811 (9308) antibodies were obtained from Cell Signaling Technology. Anti-HA antibody (3F10) was from Sigma. Anti-pRB monoclonal antibody (G3-245) was from BD Pharmingen, pRB rabbit polyclonal antibodies and E2F1 antibody (C-20) were from Santa Cruz, CAP-H2 antibody (A302-275A) was from Bethyl Laboratories. Anti-rabbit or anti-mouse IgG goat antibody conjugated to HRP was from GE Healthcare. For T-cell receptor (TCR) activation studies anti-CD3∊ (OKT3) and CD28 (CD28.2) antibodies were from BioLegend, and light chain specific goat anti-mouse IgG (115-005-174) was from Jackson ImmunoResearch Labs.

### Cell culture and stimulation

Suspension Jurkat cells were grown in DMEM enriched with 10% v/v FBS, L-glutamine, penicillin and streptomycin at 37 °C in 5% CO_2_. For SB203580 inhibitor pre-treatment, Jurkat cells (10^7^) were suspended in 5 mL enriched DMEM and SB203580 (Cell Signaling Technology) was added from 10 mM stock to a final concentration of 10 μM. Equal volume of DMSO was added to control samples. The cells were then incubated at 37 °C for 2 hours until treatment with pervanadate and calyculin (PVA/CA). Cells were centrifuged at 300 × g for 5 min at 4 °C and washed twice with PBS. After the last wash the pellet was resuspended in 5 mL enriched DMEM, and PVA/CA were added to a final concentration of 100 and 0.1 μM, respectively. Control samples received equal volumes of PBS and DMSO vehicles, respectively. Both groups were incubated at 37 °C for 20 minutes. TCR activation was achieved as previously described with modifications (21). Jurkat cells (5 × 10^6^) were resuspended in 1 mL enriched media and treated with 5 μg/mL anti-CD3∊ and CD28 antibodies, and 30 μg/mL goat anti-mouse IgG to crosslink the primary antibodies. The cells were then incubated for indicated times at 37 °C after which they were harvested.

### Protein extract preparation

All of the following lysis buffers were supplemented with 250 μM Na3VO4, 5 mM NaF, 1 mM PMSF, 5 μg/L aprotinin and leupeptin immediately before use. For whole cell extract (Ext), cells were washed twice with cold PBS and resuspended in lysis buffer containing 25 mM Tris-Cl pH 7.5, 100mM NaCl, 5% (v/v) glycerol, and 1% (v/v) NP-40 and incubated on ice for 10 min. Insoluble material was cleared by centrifugation at 12 000 × g at 4 °C for 10 min. For NE, cells were washed twice with cold PBS, resuspended in hypotonic lysis buffer containing 10 mM Tris-Cl pH 7.5, 10 mM KCl, 3 mM MgCl_2_, 1 mM EDTA and 0.05% (v/v) NP-40, and incubated on ice for 5 min. Nuclei were isolated by centrifuging the suspension at 1300 × g at 4 °C for 5 min. The pelleted nuclei were washed twice in the same buffer. The nuclei were lysed in buffer containing 20 mM Tris-Cl pH 7.5, 420 mM NaCl, 1.5 mM MgCl_2_, 0.2 mM EDTA, 25% (v/v) glycerol, 0.1% (v/v) NP-40 and incubated on ice for 10 min. Insoluble material was cleared by centrifugation at 12 000 × g at 4 °C for 10 min. For chromatin fractionation, PBS washed cells were incubated for 10 min on ice in buffer containing 10 mM Tris-Cl pH 8.0, 10 mM KCl, 1.5 mM MgCl_2_, 0.34 M sucrose, 10% (v/v) glycerol and 0.1% (v/v) Triton X-100 to lyse outer membrane. Nuclei were isolated by centrifuging the suspension at 1300 × g at 4 °C for 5 min. The pelleted nuclei were washed twice in the same buffer without Triton X-100. The nuclei were lysed in buffer containing 3 mM EDTA and 0.2 mM EGTA for 30 min on ice with occasional mixing. Nucleoplasmic fraction was removed by centrifuging the lysate at 4000 × g at 4 °C for 5 min. Remaining chromatin fraction was solubilized with DNaseI treatment at 37 °C for 10 min in buffer containing 10 mM Tris-Cl, 2.5 mM MgCl_2_, 0.5 mM CaCl_2_. Protein concentration was quantified by Bradford assay following standard procedures.

### Immunoprecipitation

Antibodies to RB or pp38 T180/Y182 were diluted 1:50 in Ext containing 1 to 2 mg total protein. The lysates were incubated at 4 °C overnight with gentle agitation. 50 μL Dynabeads Protein G (Thermo Fisher) was added and incubated at 4 °C for 2 hours with gentle agitation. A magnet was used to immobilize the beads and to discard the supernatant.

### *In vitro* kinase assay

Immunoprecipitated pp38 T180/Y182-Dynabeads complexes were used in kinase assays similar to previous reports. The complexes were resuspended in 50 μL kinase buffer containing 25 mM Tris-Cl pH 7.5, 5 mM β-glycerophosphate, 10 mM MgCl_2_, 0.25 mM Na_3_VO_4_ and 5 mM NaF. Each of the time points consisted of a 40 μL reaction containing 5 μL of immunoprecipitated kinase, 10 μg of GST-RBC, 1 mM ATP, and 50 μM SB203580 (where indicated) in kinase buffer. The reactions were prepared on ice then transferred to 37 °C to start the assay. After the given incubation times, the immunoprecipitated proteins were immobilized with a magnet and the remaining supernatant was saved.

### SDS-PAGE, immunoblotting and Coomassie staining

Protein extracts, supernatants and immunoprecipitated beads were diluted in SDS-PAGE sample buffer to a final concentration of 62.5 mM Tris-Cl pH 6.8, 10% glycerol, 2% SDS, 72.5 mM β-mercaptoethanol, and 0.005% bromophenol blue. SDS-PAGE was performed following standard procedures. After gel electrophoresis, proteins were transferred to PVDF membrane and incubated with primary antibodies diluted in 5% milk or BSA in TBS-T over night at 4 °C with gentle agitation. Membranes were washed five times in tris buffered saline-Tween 20 (TBS-T) for 5 min and incubated with horseradish peroxidase (HRP)-conjugated goat secondary antibody for one hour at room temperature. Membranes were washed again, incubated in SuperSignal WestDura (Thermo Fisher) for 5 min protected from light and developed on a ChemiDoc (Bio-rad). For Coomassie staining, gels were washed three times in water, and incubated in GelCode Blue (Thermo Fisher) over night at room temperature with gentle agitation. Stained gels were washed again twice in water. Bands were imaged on a ChemiDoc.

### Plasmids and lentiviral transduction

Briefly, pSicoR-Ef1a-mCh-Puro (Addgene #31845) was double-digested with AfeI and SmaI to replace mCherry coding sequence with that of RBLP with matching sticky ends in-frame. S838A/T841A substituted plasmid was made with QuikChange Site-directed mutagenesis kit (Agilent), following manufacturer’s recommendations and mutagenic primers TTAGTATCAATTGGTGAAGCATTCGGGGCTTCTGAGAAGTTCCAGAAA and TTTCTGGAACTTCTCAGAAGCCCCGAATGCTTCACCAATTGATACTAA. HEK293T cells at 70% confluency on 10-cm plates were transfected with 12 μg expression vector, 9 μg pMD2.G (Addgene #12259) and 3 μg psPAX2 (Addgene #12260) using Lipofectamine 3000 (Life Technologies) following manufacturer’s recommendations. Two days later, the cell media was harvested and passed through a 0.45 μm filter. Jurkat cells (10^6^) were transduced with the filtrate containing 8 μg/mL polybrene. Transduced Jurkat cells were maintained in 0.3 μg/mL puromycin.

### Chromatin sonication

Jurkat cells were washed twice in PBS, and fixed in 1% formaldehyde/PBS for 10 min at room temperature with gentle agitation. Fixation was quenched with 0.125 M glycine for 5 min at room temperature. Cells were washed twice in cold PBS and incubated for 10 min on ice in buffer containing 10 mM HEPES pH 6.5, 10 mM EDTA, 0.5 mM EGTA and 0.25% Triton X-100. Nuclei were isolated by centrifuging the suspension at 600 × g at 4 °C for 5 min. The pelleted nuclei were lysed and washed twice in buffer containing 10 mM HEPES pH 6.5, 10 mM EDTA, 0.5 mM EGTA and 200 mM NaCl. Chromatin isolated from three million cells was resuspended in 300 μL of buffer containing 50 mM Tris-Cl pH 8.0, 1 mM EDTA, 0.5 % Triton X-100 and 1% SDS, and aliquoted into 1.5 mL microtubes (Diagenode). All of the above lysis buffers were supplemented with 250 μM Na_3_VO_4_, 5 mM NaF, 1 mM PMSF, 5 μg/L aprotinin and leupeptin immediately before use. Chromatin was sonicated in Bioruptor Pico (Diagenode) at 4 °C with one to three cycles. Each cycle consisted of 15 seconds ON and 30 seconds OFF. After sonication, chromatin-protein complexes were reverse-crosslinked by adding NaCl to a final concentration of 200 mM to each sample and incubating overnight at 65 °C. RNA and protein were removed by RNase A and proteinase K digestion, respectively. DNA was isolated by phenol-chloroform extraction and ethanol precipitation, and analyzed on 3% agarose gels. DNA was stained with ethidium bromide and visualized on a Chemi Doc. High molecular weight DNA bands were quantified as a percentage of the total lane intensity with lower threshold cut-off at 100 bp using Image Lab 5.2.

### Statistical analysis

All statistical analyses were performed on Prism 8. CAPH2 chromatin loading relative to SMC1 in untransduced cells was compared using Student’s t-test. That of HA-RBLP WT or AA expressing cells was compared using two-way ANOVA and Sidak’s multiple comparisons test. For sonicated chromatin densitometry, the mean of triplicate values was compared between treatment conditions, or cell lines, as indicated by two-way ANOVA. Sidak’s multiple comparison test was used.

## DATA AVAILABILITY

All data are contained within the manuscript.

## ACKNOWLEDGEMENTS

The authors wish to thank colleagues at the London Regional Cancer Program for suggestions and encouragement in the course of this work.

## AUTHOR CONTRIBUTIONS

SJK carried out all experiments except Figure 2A, B, D, and E. JIM carried out experiments in Figure 2A, B, D, and E. SJK and FAD wrote the manuscript.

## FUNDING AND ADDITIONAL INFORMATION

SJK was funded by Cancer Research and Technology Transfer training program and Ontario Graduate Scholarship. This work was supported by a grant from the Cancer Research Society. FAD is the Wolfe Senior Fellow in Tumor Suppressor Genes at Western University.

## CONFLICT OF INTEREST

The authors declare that they have no conflicts of interest with the contents of this article.

## ABBREVIATIONS

RB: retinoblastoma tumour suppressor protein
CDK: cyclin dependent kinase
MAPK: mitogen-activated protein kinase
NHEJ: non-homologous end joining
DSB: double strand break
TCR: T-cell receptor
RBC: RB C-terminus
RBLP: RB large pocket
GST: glutathione S-transferase
PVA: pervanadate
CA: calyculin A
Ext: whole cell extract
HA: hemagglutinin

